# Genetic variants linked to neurodevelopmental disorders within the β3-β4 loop of the TRIO PH2 domain release autoinhibition of GEF2 activity

**DOI:** 10.1101/2025.04.14.648796

**Authors:** Melissa G. Carrizales, Andrew D. Boulton, Anthony J. Koleske

## Abstract

The TRIO protein contains two guanine exchange factor (GEF) domains, GEF1 and GEF2, which coordinate cytoskeletal rearrangements by activating Rho family GTPases. Rare variants that impact TRIO GEF1 function are associated with autism spectrum disorder, developmental delay, and intellectual disability, but variants are also found throughout the gene. GEF1 promotes GTP exchange on Rac1 and RhoG, while GEF2 activates RhoA. Although GEF1 and GEF2 share a common architecture, the pleckstrin homology (PH) domain in TRIO GEF1 (PH1) assists its activity, while the PH domain in GEF2 (PH2) inhibits its activation of RhoA. A series of single point variants in the unique β3-β4 loop of TRIO PH2 has been identified in patients with neurodevelopmental disorders (NDDs), but how they impact TRIO GEF2 activity is not known. Using an in vitro fluorescence-based assay to assess GEF2 exchange activity on RhoA, we demonstrate that variants within the β3-β4 loop relieve GEF2 autoinhibition. Activation of RhoA inhibits neurite outgrowth in Neuro-2A (N2A) cells. GEF2 expression in N2A cells significantly reduces neurite outgrowth, and expression of the G2211E activating GEF2 variant enhances this effect. Together, our findings reveal key interactions and structural constraints for GEF2 autoinhibition and how this mechanism is a target for disruption by NDD-associated variations.

## Introduction

Small GTPases are molecular switches that transition between inactive GDP-bound and active GTP-bound states (1,2). Active Rho GTPases play diverse roles in cytoskeletal rearrangements in various cell types (3-15). Guanine nucleotide exchange factors (GEFs) promote GTPase activation by accelerating GDP-to-GTP exchange (7,16-24). TRIO, a Rho GEF, can influence multiple Rho GTPases through its two GEF domains (GEF1 and GEF2), suggesting its participation in diverse GTPase pathways (25-33). GEF1, the most N-terminal Trio GEF domain, activates the Rac1 and RhoG GTPases, while the more C-terminal GEF2 domain activates RhoA (32,34-36). This unique aspect of TRIO is especially interesting given the opposing roles Rac1 and RhoA play in neuronal development in which Rac1 generally promotes neurite growth and stability, while RhoA destabilizes neurites and/or prevents their maturation (3,10,34,37-41). Rare predicted damaging variants in the *TRIO* gene are linked to autism spectrum disorder (ASD), intellectual disability (ID), developmental delay (DD), and schizophrenia (SCZ) and have also been observed in epilepsy (Epi) (42-57). Understanding how these variants impact TRIO function is key to elucidating the biochemical mechanisms that contribute to these disorders and potentially target these mechanisms for therapy.

While the GEF domains target different substrates, they share a common architecture, each composed of a catalytic Diffuse B cell lymphoma (Dbl) homology domain followed by a regulatory pleckstrin homology (PH) domain (25,47,58,59). Interestingly, the PH domain associated with GEF1 (PH1) promotes GEF1 activity, whereas the GEF2 PH domain (PH2) inhibits GEF2 activity (60). Bandekar et al. solved the structure of autoinhibited GEF2, uncovering a new DH-PH interface featuring a unique loop that connects the β3 and β4 sheets in PH2 (Figures 1A and 1B). This β3-β4 loop extends over the αN helix, which, based on homology comparisons with the closely related p63RhoGEF protein, is predicted to sterically block part of the RhoA binding interface (61). As such, the β3-β4 loop likely makes key contributions to suppression of GEF2 activity by PH2. Attempts to assess how removal of the β3-β4 loop impacted activity were hindered by insolubility of these mutants, so the role of this poorly conserved and disordered loop remains unknown (61). That said, the series of patient-derived variants associated with schizophrenia, bipolar disorder, and epilepsy within the β3-β4 loop of GEF2 suggests its potential in regulating either inter- or intramolecular interactions crucial for TRIO signaling.

**Figure 1.**
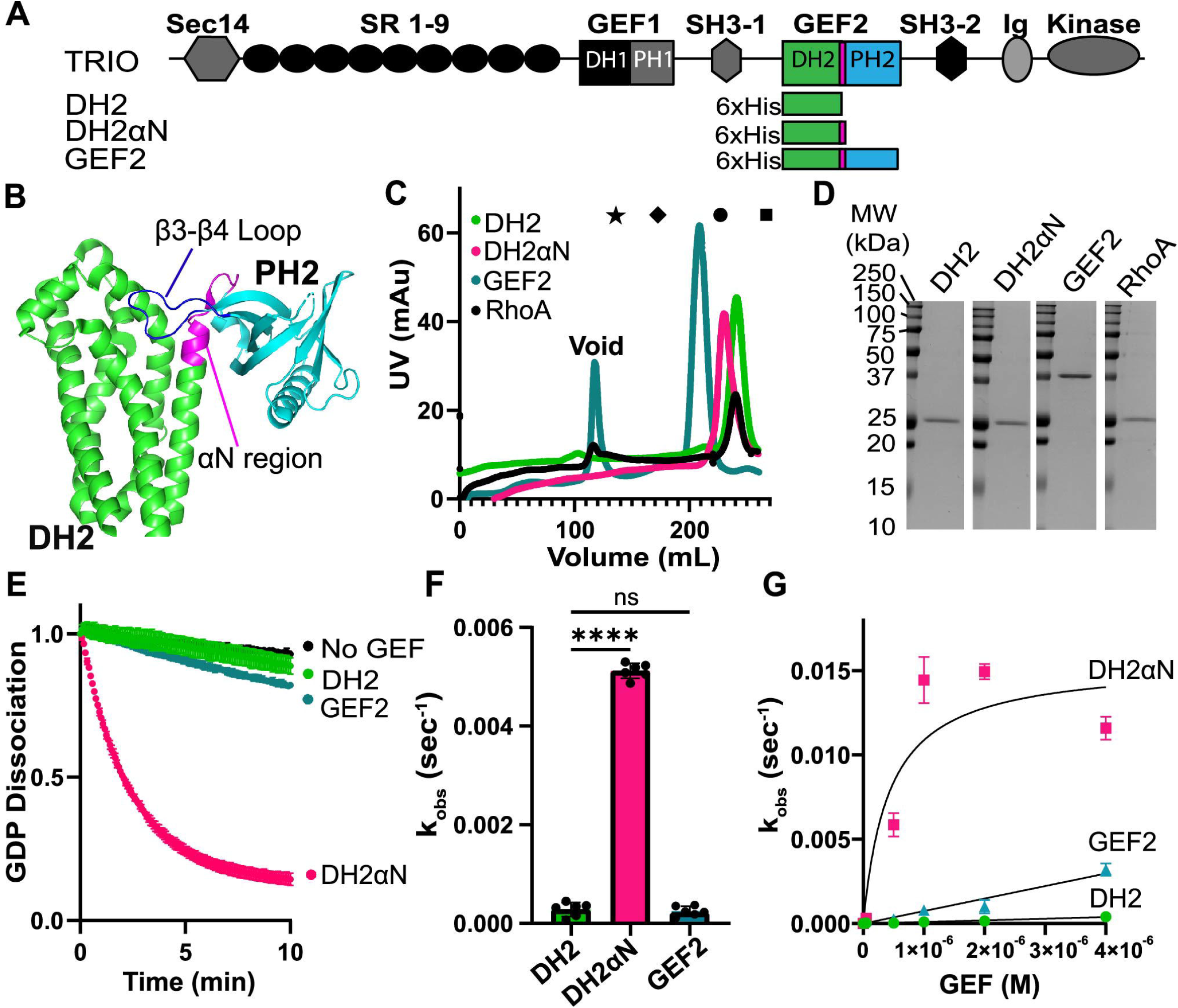
The GEF2 αN segment promotes GEF activity for RhoA. *A*, Schematic of TRIO proteins used in this study; full-length TRIO, DH2, DH2αN and GEF2; SR, spectrin repeat; DH, Dbl homology domain; PH, pleckstrin homology domain; SH3, Src homology 3 domains; Ig, Ig-like domain. *B*, GEF2 crystal structure with DH2 in green, αN in magenta, PH2 in cyan and the β3-β4 loop in dark blue (PDB 6D8Z). *C*, Elution profiles of the indicated proteins from a Superdex™ 200 Increase 10/300 GL column. Peak elution positions of commercial protein standards are represented as follows: star = 670 kDa; diamond = 150 kDa; circle = 44.3 kDa; square = 13.7 kDa. *D*, 5 µg of purified proteins were separated by SDS-PAGE and stained with Coomassie Blue to assess purity. *E*, 0.5 µM of GEF2 proteins were incubated with RhoA preloaded with 3.2 µM BODIPY-FL-GDP, and GEF activity was measured through the change in fluorescence over time. Curves represent the averages of 6 replicates ± SD. Maximal signal was normalized to 1. *F*, The first minute of activity from panel E was subjected to linear fits to extract k_obs_ for each protein; 0.5 µM DH2 had a 25-fold lower exchange rate than 0.5 µM DH2αN. Activity of the full GEF2 was similar to DH2 alone. N = 6 independent k_obs_ measurements per protein. Mean ± SD; **** p ≤ 0.0001 in a one-way ANOVA adjusted for multiple comparisons; ns = no significance. *G*, Michaelis-Menten plot of k_obs_ as a function of GEF concentration for DH2, DH2αN, and GEF2. Exponential fits to the average of 3 replicates ± SD are shown. DH2 and GEF2 plots did not achieve saturation at the concentrations tested.

Here, we test the hypothesis that the PH2 domain β3-β4 loop is a key regulator of GEF2 activation of RhoA. We present evidence that single neurodevelopmental disorder-associated missense variants within this loop can enhance GEF2 activity on RhoA and inhibit neurite development in mouse neuroblastoma cells. Together, our results clarify how the PH2 domain β3-β4 loop regulates GEF2 activity, and the impact of disease-related variants on this regulation. This mechanism may provide a candidate target for therapeutic intervention in the treatment of neurodevelopmental disorders.

## Results

### Maximal TRIO DH2 efficiency requires the αN region

Bandekar et al., showed the linker region of GEF2 includes a helix they termed the αN helix, which immediately follows DH2 (Figures 1A and 1B). This helix interacts with DH2 α helix 3 residues Glu^2069^ and Met^2146^, which are part of the predicted RhoA binding site based on structural homology to RhoA-bound p63RhoGEF (61). To determine whether the αN helix impacts the catalytic activity of the Dbl homology domain 2 (DH2), we purified TRIO DH2 alone (21 kDa) and a DH2+ αN helix residues that link the DH2 and PH2 domains (DH2αN) (22 kDa). We also purified the GEF2 unit (39 kDa), which includes the entire PH2 domain as well as the substrate RhoA GTPase (22 kDa). All proteins eluted at positions consistent with being monomers, as calibrated to commercial protein standards (Sigma-Aldrich, product #69385) (Figure 1C and 1D).

DH2 and DH2αN catalytic activity was measured using a fluorescence-based guanine nucleotide exchange assay, in which RhoA preloaded with fluorescent GDP underwent exchange with unlabeled GTP, resulting in fluorescence decay over time. Purified 0.5 µM DH2 alone catalyzed exchange of fluorescent GDP for GTP on RhoA slowly, with a first-order dissociation rate of k_obs_ = 0.000282 ± 0.000142 s^-1^, while the addition of αN accelerated GDP-dissociation by 25-fold (k_obs_ = 0.00512 ± 0.000153 s^-1^) (Figure 1E and 1F). To determine the maximum rate of reaction (V_max_), Michaelis constant (K_m_), substrate turnover (k_cat_) and catalytic efficiency (k_cat_/K_m_) of DH2 and DH2αN, we generated Michaelis-Menten plots of k_obs_ as a function of GEF concentration (Figure 1G). While a saturation curve was reached for DH2αN, saturation could not be reached within the concentration range tested for the less active DH2. The addition of αN significantly increased catalysis compared to DH2 (DH2αN k_cat_ = 0.014 ± 0.0013 s^-1^; DH2αN k_cat_/K_m_ = 3.3 × 10^4^ ± 0.619 × 10^4^ M^-1^ s^-1^). These data (summarized in Table 1) indicate that the αN segment is necessary for efficient TRIO GEF2 mediated-GDP/GTP exchange on RhoA.

### The TRIO PH2 domain decreases GEF2 catalytic activity on RhoA

While DH domains are sufficient for GEF activity, these activities can be modulated by appended PH domains. Indeed, the TRIO PH2 domain has been reported to inhibit TRIO DH2 domain activity (60). To measure how PH2 impacts catalysis of DH2αN, we generated and purified GEF2 which includes the full DH2αN+PH2 unit (Figure 1A-D). GEF2 showed a 22-fold decrease in catalytic rate (GEF2_WT_ k_obs_ = 0.000222 ± 0.000106 s^-^1) relative to DH2αN (Figure 1E, 1F and 1G). Like DH2, GEF2 did not saturate activity in our range of concentration. These observations indicate that the PH2 domain reduces DH2αN-mediated GDP/GTP exchange on RhoA.

### NDD-associated variants within the β3-β4 loop impact GEF2 catalytic activity

We explored variants related to NDDs from the online database Exome Sequencing Meta-analysis (SCHEMA, https://schema.broadinstitute.org/). We found six single-substitution variants near the αN region of GEF2 are located within the β3-β4 loop of GEF2 in individuals with bipolar disorder, schizophrenia, and epilepsy (Figure 2A). We focused on these variants going forward: P2200S, L2201H, D2202G, K2204Q, P2210L and G2211E.

**Figure 2.**
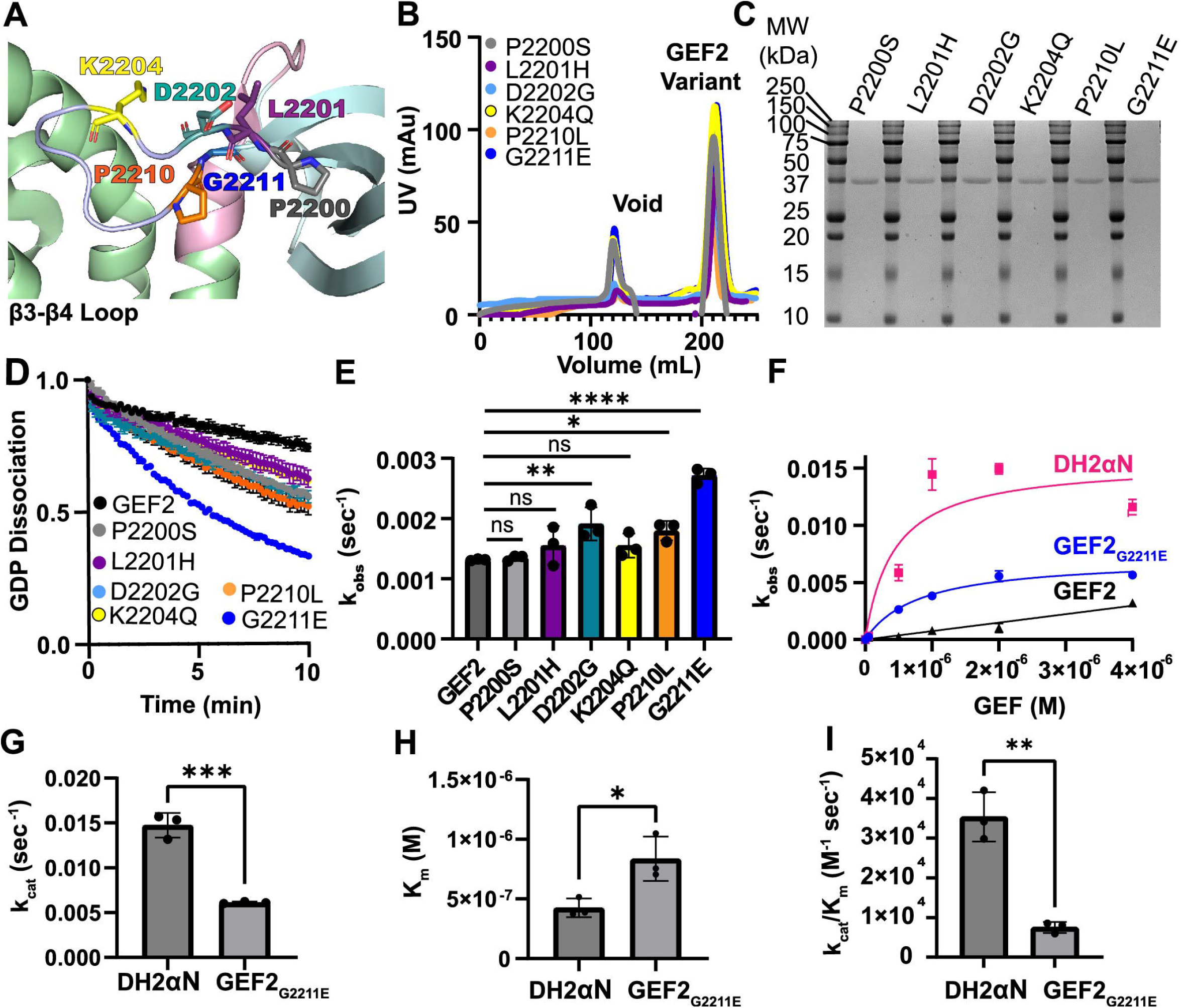
NDD-associated variants in the PH2 β3-β4 loop release GEF2 autoinhibition. *A*, GEF2 crystal structure (PDB 6D8Z) showing the position of NDD-associated variants within the β3-β4 loop; DH2 domain in green; αN in pink; PH2 domain in cyan; Target residues for variation colored as follows: P2200 in gray; L2201 in purple; D2202 in teal; K2204 in yellow; P2210 in orange and G2211 in blue. *B*, Elution profiles of the indicated proteins. *C*, 5 µg of purified components were separated by SDS-PAGE and stained with Coomassie Blue to assess purity. *D*, 0.5 µM of GEF2 variant proteins were incubated with RhoA preloaded BODIPY-FL-GDP, and GEF activity was measured by monitoring fluorescence over time. Curves represent the averages of 3 replicates ± SD. *E,* GEF2 P2200S (BP), L2201H (SCZ), and K2204Q (SCZ) did not impact exchange rate relative to GEF2, while D2202G (SCZ), and P2210L and G2211E (both Epi) increased GEF2 activity. N = 3 independent k_obs_ measurements per variant. Bars represent mean ± SD; **** = *p* ≤ 0.0001 in a one-way ANOVA adjusted for multiple comparisons. *F*, Michaelis-Menten plot of k_obs_ as a function of GEF concentration for GEF2 and select variants. N= 3 replicates ± SD. GEF2 did not saturate at the concentrations tested. The data for DH2αN and GEF2 are also present in Figure 1. *G*, Significant differences were observed for substrate turnover (k_cat_) between DH2αN and the G2211E mutation. Bars represent mean ± SD. N = 3 independent k_cat_ measurements per group. \*\*\**p* ≤ 0.001 in a one-way ANOVA adjusted for multiple comparisons. *H*, Significant differences were observed for substrate recognition (K_m_) between DH2αN and the G2211E mutation. N = 3 independent K_m_ measurements per group. Bars represent mean ± SD; \**p* ≤ 0.05 in a one-way ANOVA adjusted for multiple comparisons. *I*, Significant differences in catalytic efficiency (k_cat_/K_m_) were observed between DH2αN and the G2211E variant. N = 3 independent k_cat_/K_m_ measurements per group. Bars represent mean ± SD; \*\**p* ≤ 0.01 in a one-way ANOVA adjusted for multiple comparisons.

We created GEF2 expression constructs containing single NDD-associated point mutations and measured impacts to nucleotide exchange on RhoA (Figures 2B and 2C). The D2202G (SCZ), and P2210L and G2211E (both Epi) increased k_obs_ by 1.5-fold,1.4-fold and 2-fold over WT GEF2, respectively (GEF2_WT_ k_obs_ = 0.0013 ± 0.00017 s^-^1, GEF2_D2202G_ k_obs_ = 0.0019 ± 0.00027 s^-1^, GEF2_P2210L_ k_obs_ = 0.0018 ± 0.00016 s^-1^, GEF2_G2211E_ k_obs_ = 0.0027 ± 0.00012 s^-1^) (Figures 2D and 2E).

Maximum catalytic rate could be saturated under the concentration range tested for GEF2_G2211E_ (Figure 2F). We found that the K_m_ was increased approximately 2-fold as compared to active DH2αN (GEF2_G2211E_ K_m_ = 8.35 × 10^−7^ ± 1.85 × 10^−7^ M ; DH2αN K_m_ = 4.25 × 10^−7^ ± 7.82 × 10^−8^ M) while GEF2_G2211E_ was 4.7-fold slower than DH2αN in catalytic rate (GEF2_G2211E_ k_cat_ = 0.006 ± 0.00012 s^-1^, k_cat_/K_m_ = 0.752 x10^4^ ± 0.137 × 10^4^ M^-1^ s^-1^) (Figures 2G-I). These findings are summarized in Table 1 and provide evidence that missense substitutions in the β3-β4 loop of PH2 can relieve autoinhibition.

### G2211E promotes inhibition of N2A neurite outgrowth by GEF2

RhoA activity, through its activation of ROCK kinase, formins, and other effectors, regulates actin assembly and actomyosin contractility. Attenuation of RhoA activity is required for neurite sprouting formation in Neuro-2A (N2A) neuroblastoma cells following serum withdrawal (63). To test how their relative GEF2 activities impacted neurite outgrowth, we expressed GFP fusion proteins, DH2αN-GFP, GEF2-GFP, GEF2_G2211E_-GFP, and a GEF2 mutant we engineered to have reduced catalytic activity, termed GEF2_N2143A/D2144A_ (Figure 3A), in N2A cells at similar levels (Figures 3B and 3C) and ensured transfection conditions did not alter endogenous RhoA expression (Figures 3B and 3E). We measured neurites at 48 hrs after serum starvation (64) (Figures 3F-K).

**Figure 3.**
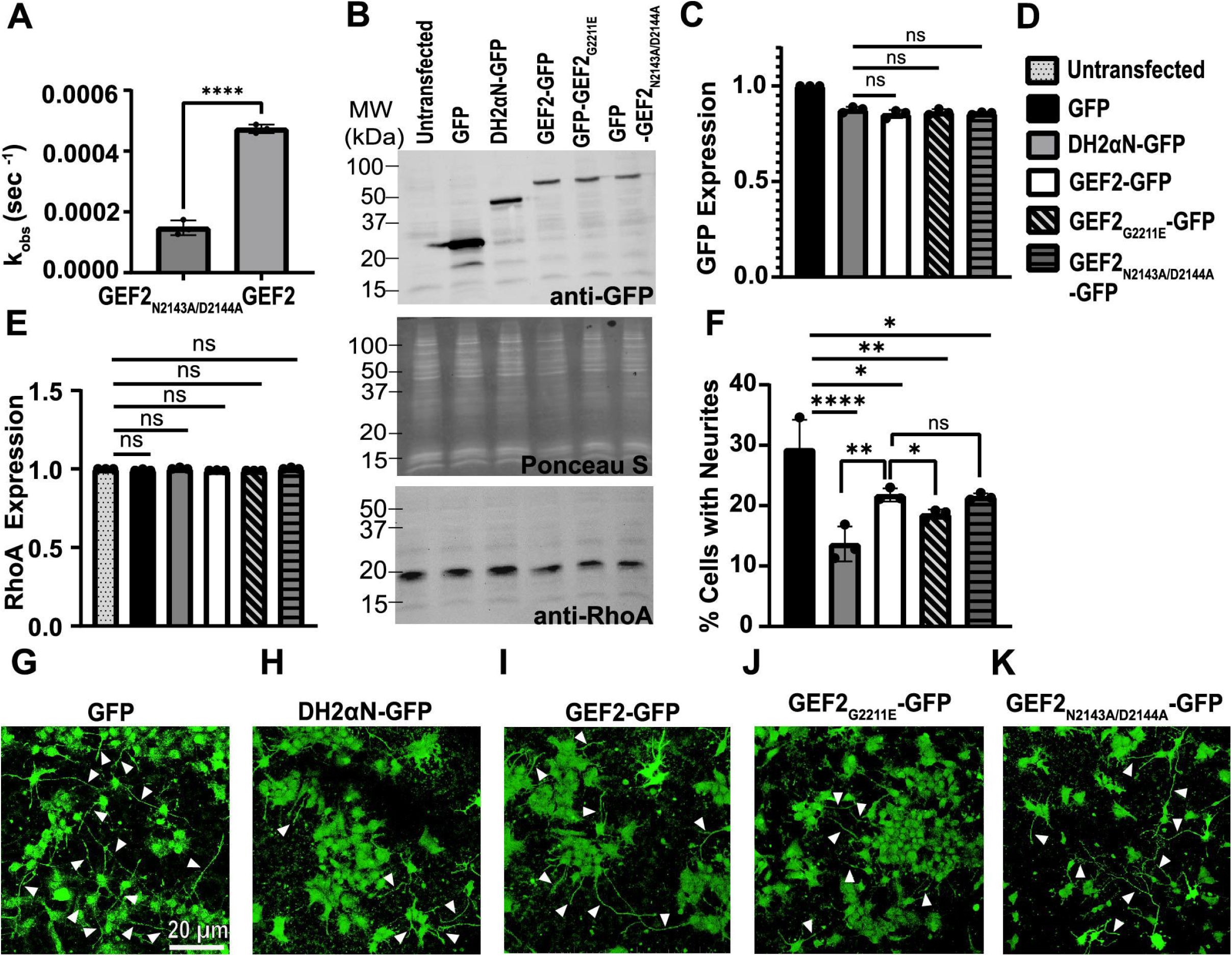
Activated GEF2 reduces neuritogenesis in N2A Cells. *A*, GEF2_N2143A/D2144A_ and GEF2 proteins were tested with RhoA preloaded with BODIPY-FL-GDP, and GEF activity was measured by monitoring fluorescence over time. N = 3 independent k_obs_ measurements per protein; Mean ± SD; ****p ≤ 0.0001 in a one-way ANOVA adjusted for multiple comparisons. *B, and C*, Expression of each construct was controlled for equal expression; pBlueScriptII SK+ DNA was used to maintain DNA levels between all transfections when necessary; N = 3 independent transfections per group; Bars represent mean ± SD; ns = no significance in a one-way ANOVA adjusted for multiple comparisons; GFP signal consistently remained slightly higher; S Ponceau is the loading control. *D*, Legend for panels C, E and F. *E*, Endogenous RhoA expression was not impacted by transfections; N = 3 independent transfections per group; Bars represent mean ± SD; ns = no significance in a one-way ANOVA adjusted for multiple comparisons. *F*, Percent of cells with neurites from N2A cells; mean ± SD, N = 3 transfections; ****p ≤ 0.0001 in a one-way ANOVA adjusted for multiple comparisons. *G–K*, Representative images of N2A cells expressing GFP fusion proteins plated on poly-L-ornithine and laminin-coated cover slips for GFP (*G*), DH2αN-GFP (*H*), GEF2-GFP (*I*), GEF2_G2211E_-GFP (*J*), and GEF2_N2143A/D2144A_-GFP (*K*); white arrows point to a neurite.

Even though GEF2 is autoinhibited by its PH2 domain, some basal activity remains, and we see this reflected as a 7.7% decrease in neurite outgrowth (GFP_control_ = 29.45 ± 4.7%; GEF2-GFP = 21.8 ± 1.06%). As expected, cells expressing catalytically active proteins, DH2αN and GEF2_G2211E_, had higher reductions of neurite outgrowth by 15.78 % and 10.85 % respectively (DH2αN-GFP = 13.67 ± 2.8%; GEF2_G2211E_-GFP = 18.6 ± 0.79%). In addition, the reduced activity GEF2_N2143A/D2144A_-GFP inhibited neurite outgrowth to an extent similar to GEF2 (GEF2_N2143A/D2144A_ = 21.5 ± 0.56%). These experiments strongly suggest that the impact of the G2211E variant on GEF2 activity is sufficient to impact the magnitude of a RhoA-dependent process in N2A cells.

## Discussion

Our study reveals fundamental principles and determinants of the catalytic efficiency and regulation of GEF2, a guanine nucleotide exchange factor (GEF) that plays a pivotal role in regulating RhoA activity during neuronal differentiation. The GEF2 domain consists of a DH and PH domain as defined by homology to other DH and PH domains, connected by a unique sequence of residues, including the αN helix. This helix follows the DH2 domain, extending the C-terminal helical structure beyond the canonical DH2 domain. Homology studies with the closely related protein p63RhoGEF suggest that this C-terminal extension contributes to a platform for interactions with RhoA (61). Our findings demonstrate that the inclusion of the αN helix significantly enhances GDP/GTP exchange on RhoA, increasing both the dissociation rate and overall catalytic efficiency, highlighting the essential role of the αN helix in catalysis by GEF2.

We also demonstrate that the TRIO PH2 domain inhibits the catalytic activity of DH2αN. The presence of PH2 reduces the GDP/GTP exchange rate on RhoA by 22-fold, suggesting that PH2 acts as an autoinhibitory regulator. This finding aligns with previous reports showing that PH domains modulate GEF activity (60), and suggests PH2 as a mechanism to control the timing and extent of RhoA activation during neuronal development. The unique role of the PH2 domain in GEF2 regulating RhoA activation is supported by evidence that swapping the PH domain of GEF2 with that of GEF1 increases RhoA exchange activity and reduces Rac1 exchange (60). This inhibitory role may be conserved in some RhoA GEFs, as seen in p63RhoGEF, which carries a GEF domain similar to TRIO GEF2. Previous studies also identified p69RhoGEF as a key Gαq effector crucial for behaviors such as locomotion and egg laying in *C. elegans*, with its function relying on a conserved C-terminal extension of the PH domain, shared with TRIO PH2 (65). While direct interactors of TRIO PH2 are not yet known, it is suggested that PH domains bind lipids, particularly phosphoinositides, which are key components of biological membranes (66-68). This implies that TRIO localization and its GEF1 and GEF2 activities may be regulated through its two PH domains, potentially within specialized nanodomains in the neuronal membrane, such as PIP3-rich regions in growth cone membranes (69). Increasing evidence reveals how individual PH domains regulate GEF activity in distinct ways, highlighting the diversity in how PH domains control Rho family GTPase signaling.

Variants in TRIO are associated with neurodevelopmental disorders. In TRIO’s GEF2 unit, we identified several disorder-associated variants in the β3-β4 loop of PH2, near the αN helix. Introduction of these variants, found in individuals with bipolar disorder, schizophrenia, and epilepsy, into GEF2 altered nucleotide exchange rates on RhoA, with some mutations significantly increasing the catalytic activity of GEF2. In particular, the G2211E variant exhibited a 2-fold increase in k_obs_, indicating that variants in this region at least partly relieves the autoinhibition imposed by the PH2 domain.

Finally, we explored whether GEF2 can encourage inhibition of neurite outgrowth, a RhoA dependent process, by assessing neurite extension in N2A cells. The overexpression of various GEF2 constructs, including the G2211E variant, led to reduced neurite extension, with the G2211E variant showing a significant reduction of 10.82%. This suggests that disease-associated mutations in GEF2 not only affect its biochemical activity but also have functional consequences on neuronal morphogenesis. To understand the molecular basis for how G2211E variant might impact GEF2 catalysis, we utilized AlphaFold 3 to predict its impact on structure (Figures 4A-C). Our analysis revealed that the G2211E variant is predicted to shift the β3-β4 loop, which exposes the αN helix residues for enhanced interactions with RhoA (61). This structural change could explain the increased catalytic activity observed with the G2211E variant, providing insight into how this mutation might disrupt GEF2 regulation and contribute to neurodevelopmental disorders.

**Figure 4.**
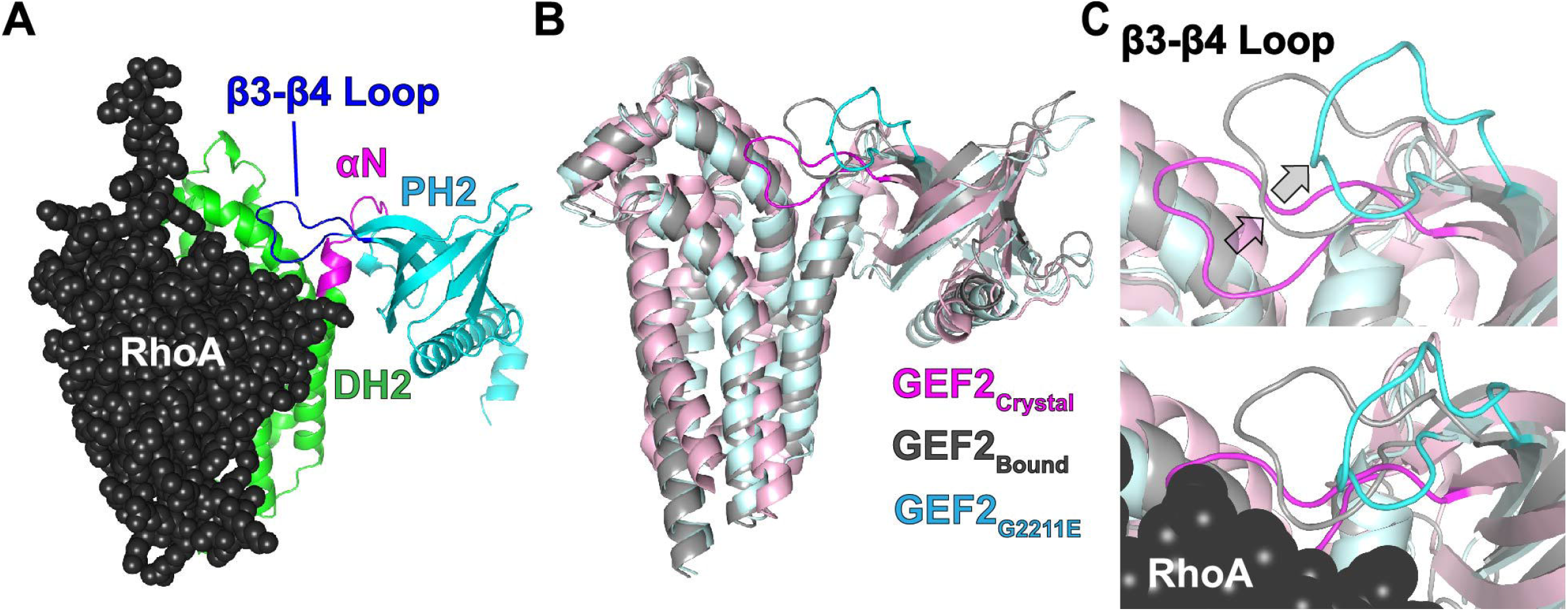
Alpha Fold 3 predicts a structural shift of the β3-β4 loop in PH2. *A*, The GEF2 crystal structure (PDB 6D8Z) was modeled bound to RhoA in Alpha Fold 3; pLDDT = 90.4; pTM = 0.779. *B*, The unbound, autoinhibited crystal structure (GEF2_crystal_) was aligned to the predicted GEF2:RhoA complex (GEF2_bound_) and the predicted structure for the G2211E mutation (GEF2_G2211E_); RhoA was hidden for clarity. *C*, Focused view of the β3-β4 loop from the same alignment as B; RhoA is hidden for clarity in the top panel.

Taken together, our findings support a critical role for GEF2 in regulating RhoA during cellular differentiation and suggest that disruptions in its function, through mutations or altered regulatory mechanisms, may contribute to neurodevelopmental disorders. Further studies will be necessary to investigate the broader implications of these mutations in vivo and to explore potential therapeutic approaches for modulating GEF2 activity in disease contexts.

## Experimental procedures

### Cloning of expression constructs and protein purification

Human TRIO GEF2 and RhoA constructs were created and affinity purified from bacterial cells as described (62). Eluted protein was further purified over an S200 Increase column into assay buffer (50 mM Hepes pH 7.25, 150 mM KCl, 5% glycerol, 0.01% Triton, 1 mM DTT) and flash frozen for storage at -80C. Site-directed mutagenesis was used for point mutations of GEF2. All constructs were confirmed by DNA sequencing. Primers used for cloning are detailed in Supporting Information.

### GEF2 nucleotide exchange assays

RhoA was loaded with 3.2 µM BODIPY-FL-GDP (Invitrogen) in assay buffer with 2 mM EDTA and incubated for 1 h at room temperature and the loading reaction was halted by adding 5 mM MgCl_2_. Each 30 µl reaction was loaded into 384-well, glass bottom black wall plates (Corning). Prior to initiating the reaction, 0-4 µM TRIO GEF2 proteins were combined with 4 mM GTP in a final volume of 10 µl. Exchange reactions were initiated by adding the 10 µL of the GEF2/GTP mix into the 30 µl in the plate wells for a total reaction mixture of 40 µl. Real-time fluorescence data was measured every 10 s for 10 min monitoring BODIPY-FL fluorescence by excitation at 488 nm and emission at 535 nm, as described in Blaise et al (62). Plots of fluorescence decay *versus* time (in seconds) were generated in GraphPad Prism 10 and the rate constant, k_obs_, was determined from the initial slope of the reaction using Equation A, where Y is the signal for GDP dissociation, and X is time in seconds.

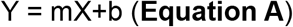

Measurements of GEF2 catalytic efficiency were conducted similarly with RhoA concentration fixed at 0.5 µM and increasing concentrations of GEF proteins (0.05 µM, 0.5 µM, 1 µM, 2 µM, and 4 µM). At least three replicates per GEF concentration were performed. Michaelis-Menten plots were created by plotting k_obs_ value*s versus* GEF concentration. From this plot, the Michaelis-Menten constant (K_m_) was extracted using Equation B in GraphPad Prism 10, where K_m_ is in M.

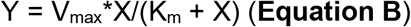

The catalytic constant, k_cat_, values were extracted using **Equation C** below in GraphPad Prism 10, where E_t_ is the GEF concentration. Catalytic specificity was determined by the ratio of k_cat_/K_m_.

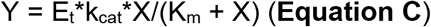

### Transfection of Neuroblastoma Cells

N2A cells were plated on 6 cm plates with DMEM containing 10% FBS and transfected with 6 µg of plasmid DNAs per plate using Lipofectamine 2000 (Invitrogen). After 48 hours cells were used for immunoblot analysis or scored for neurites after serum starvation with DMEM containing 0.5% FBS. Cells that had neurites greater than twice the cell body length were scored as cells with neurites by a reviewer blinded to transfection conditions. 0.5 µg of N1-mCherry was co-transfected to visualize transfection efficiency. pBlueScript II SK+ DNA was used to maintain equal DNA levels during transfection when necessary. Images were obtained at 20 X magnification using the Dragonfly 630 confocal microscope (Andor).

### Immunoblot Analysis

5 µg of cell lysate containing each GFP fusion protein were resolved by SDS-PAGE and transferred to nitrocellulose membranes. Protein detection was performed using one of the following: (1) Ponceau S staining, (2) anti-GFP antibody (Invitrogen, PA1-28664), or (3) anti-RhoA antibody (Invitrogen, MA1-134). Blots were quantified using ImageJ. Protein band intensities were normalized to total protein content as determined by Ponceau S staining. For comparative analysis, chemiluminescent signals were normalized to the GFP control.

### Protein structure predictions

AlphaFold 3 was used to predict the structure of human GEF2 (residues 1960–2292) in complex with human RhoA and to model the structural impact of the G2211E mutation. Predicted models were aligned to the GEF2 crystal structure (PDB: 6D8Z) using PyMOL v2.6 for comparative analysis.

### Rigor and Statistical Methods

At least three individual replicates for nucleotide exchange assays per GEF protein and concentration were conducted, and results show means ± SD. Rate of decay was extracted via a simple linear regression fit to the initial slopes of decay curves in GraphPad Prism 10. A one-way ANOVA was used to determine statistical significance of rate constants between GEF2, DH2, DH2αN and GEF2 variants (two-tailed p-value < 0.05) and adjusted using Dunnett’s multiple comparisons test.

For cell-based assays, three individual transfections were conducted for DH2αN, GEF2 and GEF2 variants, and results show means ± SD. One-way ANOVA was used to determine statistical significance of neurite formation between DH2αN, GEF2 and GEF2 variants (two-tailed p-value < 0.05) and adjusted using Dunnett’s multiple comparisons test.

## Supporting information

Primers

## Data Availability

Data available upon request. Contact for more information.

## Supporting Information

This article contains supporting information.

## Acknowledgements

We thank Xianyun Ye and Xiaoyuan Li for expert technical support. We also thank Robert Niescier (R. N.) and Shengyu Zhao (S. Z.) for review and editing of this manuscript.

## Author Contributions

M. G. C, and A. J. K. conceptualization; M. G. C., A. J. K., and A. D. B. methodology; M. G. C. formal analysis; M. G. C. investigation; M. G. C., and A. J. K. writing–original draft; M. G. C. visualization; M. G. C., and A. J. K. funding acquisition; M. G. C., and A. J. K. project administration; A. D. B., A. J. K., R. N., and S. Z. writing– review and editing; A. J. K. supervision

## Funding and additional information

This work was supported by NIH grants F31NS131038 (M.G.C.) and R01MH132685 and R01MH133562 (A.J.K.)

## Abbreviations

DH1: Dbl homology domain 1
DH2: Dbl homology domain 2
GEF: guanine exchange factor
NDD: neurodevelopmental disorder
PH1: pleckstrin homology domain 1
PH2: pleckstrin homology domain 2
ASD: autism spectrum disorder
SCZ: schizophrenia
Epi: epilepsies
ID: intellectual disability
DD: developmental delay

