## Supplementary material for "Genetic variants linked to neurodevelopmental disorders within the β3-β4 loop of the TRIO PH2 domain release autoinhibition of GEF2 activity": Table 1

| GEF Protein | $k_{obs}$ ($s^{-1}$) | Fold Activation over GEF2 for $k_{obs}$ ($s^{-1}$) | $k_{cat}$ ($s^{-1}$) | $K_m$ (M) | $k_{cat}/K_m$ ($M^{-1}$ $s^{-1}$) |
|---|---|---|---|---|---|
| DH2 | 0.000282 ± 0.000142 | NA | | | |
| DH2αN | 0.00512 ± 0.000153 | 22 | 0.014 ± 0.0013 | $4.25 \times 10^{-7} \pm 7.82 \times 10^{-8}$ | $3.3 \times 10^{4} \pm 0.619 \times 10^{4}$ |
| $GEF2_{WT}$ | 0.000222 ± 0.000106 | NA | | | |
| $GEF2_{WT}$ | 0.0013 ± 0.00017 | NA | | | |
| $GEF2_{P2200S}$ | 0.001358 ± 0.000044 | NA | | | |
| $GEF2_{L2201H}$ | 0.00155 ± 0.0003201 | NA | | | |
| $GEF2_{D2202G}$ | 0.0019 ± 0.00027 | 1.5 | | | |
| $GEF2_{K2204Q}$ | 0.00155 ± 0.00019 | NA | | | |
| $GEF2_{P2210L}$ | 0.0018 ± 0.00016 | 1.4 | | | |
| $GEF2_{G2211E}$ | 0.0027 ± 0.00012 | 2 | 0.006 ± 0.00012 | $8.35 \times 10^{-7} \pm 1.85 \times 10^{-7}$ | $0.752 \times 10^{4} \pm 0.137 \times 10^{4}$ |
| $GEF2_{N2143A/D2144A}$ | 0.000148 ± 0.000056 | NA | | | |

**Table 1. Summary of in vitro GEF2 nucleotide exchange assays**. 0.5 µM RhoA was loaded with BODIPY-FL-GDP in assay buffer, and the reaction was stopped with 5 mM $MgCl_2$ after a 1 hr incubation. Prior to initiating exchange, 0–4 µM TRIO GEF proteins were mixed with 4 mM GTP. Reactions were started by adding the GEF2/GTP mix to the loaded RhoA. Fluorescence was recorded every 10 s for 10 min to monitor BODIPY-FL-GDP release, as described in Blaise et al. (62). Fluorescence decay curves were analyzed in GraphPad Prism 10, and the rate constant ($k_{obs}$) was calculated from the initial slope. Measurements of GEF2 catalytic efficiency were conducted similarly with RhoA concentration fixed at 0.5 µM and increasing concentrations of GEF proteins. Michaelis-Menten plots were created by plotting $k_{obs}$ values versus GEF concentration. From this plot, the Michaelis-Menten constant ($K_m$) and the catalytic constant, $k_{cat}$, were extracted.
