## Supplementary material for "Genetic variants linked to neurodevelopmental disorders within the β3-β4 loop of the TRIO PH2 domain release autoinhibition of GEF2 activity": Primers

| construct | nt change | primer seq fwd (5'-3') | primer seq rev (5'-3') | vector | Comments |
|---|---|---|---|---|---|
| DH2 | NA | GATC**gaattc**GAAGAAAGGAAATCCAGCTCTTTAAAG | GATC**gcggccgc**TCACATGTCGTTGCACCGCCTGGG | pET-His-TT | lower case and bold = restriction site |
| DH2αN | NA | GATC**gaattc**GAAGAAAGGAAATCCAGCTCTTTAAAG | GATC**gcggccgc**TCACCCGTCGAATCCTTGCAGC | pET-His-TT | lower case and bold = restriction site |
| GEF2 | NA | GATC**gaattc**GAAGAAAGGAAATCCAGCTCTTTAAAG[illegible] | GATC**gcggccgc**TCATTCTAAAATTTGGTTGATTTCATGG | pET-His-TT | lower case and bold = restriction site |
| GEF2$_{P2200S}$ | c6598t | GATCGTCATATTCAGCGAA**t**CACTTGATAAAAAGAA[illegible] | CTTTTTATCAAGTG**a**TTCGCTGAATATGACGATCTGC | pET-His-TT | lower case and bold = nucleotide change |
| GEF2$_{L2201H}$ | t6602a | CAGCGAACCAC**a**TGATAAAAAGAAGGGCTTCTC | GAGAAGCCCTTCTTTTTATCA**t**GTGGTTCGCTG | pET-His-TT | lower case and bold = nucleotide change |
| GEF2$_{D2202G}$ | a6605g | CAGCGAACCACTTG**g**TAAAAAGAAGGGCTTC | GAGAAGCCCTTCTTTTTA**c**CAAGTGGTTCGCTG | pET-His-TT | lower case and bold = nucleotide change |
| GEF2$_{K2204Q}$ | a6610c | CCACTTGATAAA**c**AGAAGGGCTTCTCCATGCCG | CGGCATGGAGAAGCCCTTCT**g**TTTATCAAGTGG | pET-His-TT | lower case and bold = nucleotide change |
| GEF2$_{P2210L}$ | c6629t | GAAGGGCTTCTCCATGC**t**GGGATTCCTGTTTAAG | CTTAAACAGGAATCCC**a**GCATGGAGAAGCCCTTC | pET-His-TT | lower case and bold = nucleotide change |
| GEF2$_{G2211E}$ | g6632a | GGCTTCTCCATGCCGG**a**ATTCCTGTTTAAGAACAG | CTGTTCTTAAACAGGAAT**t**CCGGCATGGAGAAGCC | pET-His-TT | lower case and bold = nucleotide change |
